# Spatial patterns in functional brain connectomes reflect acute and chronic cannabis use

**DOI:** 10.1101/2021.03.08.434333

**Authors:** JG Ramaekers, NL Mason, SW Toennes, EL Theunissen, E Amico

## Abstract

Resting state fMRI has been employed to identify alterations in functional connectivity within or between brain regions following acute and chronic exposure to Δ9-tetrahydrocannabinol (THC), the psychoactive component in cannabis. Most studies focused a priori on a limited number of local brain areas or circuits, without considering the impact of cannabis on wholebrain network organization. The present study attempted to identify changes in the wholebrain human functional connectome as assessed with ultra-high field (7T) resting state scans of occasional (N=12) and chronic cannabis users (N=14) during placebo and following vaporization of cannabis. Two distinct data-driven methodologies, i.e. network-based statistics (NBS) and connICA, were used to identify changes in functional connectomes associated with acute cannabis intoxication and chronic cannabis use. Both methodologies revealed a broad state of hyperconnectivity within the entire range of major brain networks in chronic cannabis users compared to occasional cannabis users, which might be reflective of an adaptive network reorganization following prolonged cannabis exposure. The connICA methodology also extracted a distinct spatial connectivity pattern of hypoconnectivity involving the dorsal attention, limbic, subcortical and cerebellum networks and of hyperconnectivity between the default mode and ventral attention network, that was associated with the feeling of subjective high during THC intoxication across both user groups. Whole-brain network approaches identified spatial patterns in functional brain connectomes that distinguished acute from chronic cannabis use, and offer an important utility for probing the interplay between short and long-term alterations in functional brain dynamics when progressing from occasional to chronic use of cannabis.

## Introduction

The endocannabinoid system has been implicated to play a modulatory role in cognition and motor function, neuroprotection, nociception, synaptic plasticity and inflammation ^1^. Cannabinoid type 1 (CB1) receptors are widely expressed in the brain at presynaptic terminals that are activated by endocannabinoids, a group of retrograde neurotransmitters that include anandamide and 2-arachidonoylglycerol ^2^. Activation of CB1 receptors leads to suppression of glutamate and GABA release from the presynaptic terminal and modulates a wider range of neurotransmitter circuits of which they are part ^3,4^. CB1 receptor activation is also thought to be responsible for the disruptive effects on human brain function, cognition and psychomotor performance caused by exogenous cannabinoids such as Δ9-tetrahydrocannabinol (THC), the psychoactive ingredient of cannabis ^5–12^. A number of human studies have confirmed that psychoactive and physiological effects of cannabis can be successfully blocked or attenuated by the coadministration of a CB1 antagonist ^13–16^.

In addition to the acute psychoactive effects of cannabis, studies have also demonstrated residual cognitive deficits in chronic cannabis users ^17–19^. Typically, such deficits decrease during abstinence and do not persist beyond 4-5 weeks ^17,20^. Likewise, studies have shown that cortical CB1 receptors become downregulated with years of cannabis use, but quickly start to recover within days and return to control levels within 4 weeks of abstinence ^21–23^. These findings suggest that changes in CB1 receptor signaling contribute to the development of cognitive deficits resulting from chronic exposure to cannabis and that recoveries of CB1 receptors and cognitive deficits observed during cannabis abstinence are related ^24^.

Recent advances in functional magnetic resonance imaging (fMRI) have allowed researchers to investigate neuronal-related temporal fluctuations in the activity of different areas in the brain. The study of the pairwise correlated/anti-correlated activity between different brain regions has become popularly known in the research community as “functional connectivity” ^25^. Functional connectivity measures have been employed to identify brain areas that underlie acute and chronic cannabis effects on cognitive function. Overall, these analyses have shown that acute THC intoxication causes reductions in functional connectivity within the mesocorticolimbic circuit and the salience network, and that these changes are associated with decrements in cognitive function or increments in psychotomimetic symptoms ^10,11,26–30^. Chronic use of cannabis has also been associated with a range of functional connectivity alterations that can be measured during abstinence. Hypoconnectivity in corticostriatal circuits was associated with anhedonia ^31,32^, whereas increased functional connectivity in the posterior cingulate cortex alongside reduced functional connectivity in the hippocampus was associated with memory impairment in chronic cannabis users as compared to controls ^33^. Chronic cannabis users also displayed increased functional connectivity between frontal brain areas and subcortical regions ^34,35^ that have been associated to impulsive behavior ^35^ and mood ^36^. Likewise subcortical hyperconnectivity has been reported in cannabis dependence ^37^.

Most functional connectivity studies in cannabis users focused *a priori* on a limited number of local brain areas to define acute and residual effects of THC on brain function. Only one study so far has attempted to define patterns of whole-brain functional connectivity within and between the entire range of available brain areas to characterize cannabis induced states ^38^. That study revealed an association between cannabis intoxication and a specific pattern of functional connectivity alterations within and between auditory and somato-motor cortices that were anti-correlated to alterations in subcortical structures and the cerebellum. The relevance of looking at the entire human functional connectome is that it might allow delineation of comprehensive patterns of functional connectivity in response to acute as well as chronic exposure to cannabis. Recent advances in functional neuroimaging have provided new tools to measure these connections in disease states, e.g. neurological disorders and alcohol abuse ^39–41^, by studying the brain as a functional network (also called “functional connectome” or “functional connectivity matrix”) and by extracting connectivity patterns relevant to the disease at hand ^42^. Here we hypothesized that such patterns of connectivity might fluctuate as a function of cannabis use history and transiently change during cannabis exposure. In turn, this might serve as important markers of brain function, particularly when their variability is associated with alterations in cognitive and behavioral variables that affect real-world functions of cannabis users.

The present study therefore attempted to identify changes in the whole-brain human functional connectome of occasional and chronic cannabis users during a placebo treatment and following vaporization of cannabis. We used two distinct data-driven methodologies to identify changes in functional connectomes. The first, Network-Based Statistics (NBS), is a common procedure to make statistical inferences on functional connectomes ^43^. The second, connICA ^44^, uses independent component analysis in the connectivity domain to extract patterns of connectivity that are associated with clinical characteristics such as, in the case of this work, cannabis use history and level of cannabis intoxication. We expected that this first functional connectome-based investigation would offer unique insights into the alterations of human brain networks’ connectivity following acute and chronic cannabis use.

## Methods

### Participants, design and procedures

Participants were recruited through advertisements around Maastricht University. Inclusion criteria were: age, 18-40 years; occasional cannabis use for the occasional group, ranging between 1 time a month and 3 times a week for the past year; chronic cannabis use for the chronic group, using at least 4 times a week for the past year; body mass index between 18 and 28 kg/m2; and written informed consent. Exclusion criteria were: history of drug abuse (other than the use of cannabis) or addiction; pregnancy or lactation; health issues including hypertension (diastolic >90 and systolic >140), cardiac dysfunction, and liver dysfunction; current or history of endocrine, neurological, psychiatric disorders; use of psychotropic medication; previous experience of serious side effects to cannabis; and MRI contraindications. Before inclusion, subjects were screened and examined by a study physician, who checked for general health, conducted a resting ECG, and took blood and urine samples in which hematology, clinical chemistry, urine, and virology analyses were conducted. Participant demographic data can be found in Table S1. Overall, demographics did not differ between groups, except for their frequency of cannabis use.

The study was conducted according to a double-blind, placebo-controlled, mixed cross-over design in occasional (N=14) and chronic cannabis users (N=12). Each participant received cannabis placebo and cannabis (300 μg/kg THC) on separate days, separated by a minimum wash-out period of 7 days. Medical cannabis (Bedrobinol; 13.5% THC) was obtained from Bedrocan, the Netherlands. Treatment orders were randomly assigned to participants. Cannabis and cannabis placebo were administered through a Volcano vaporizer (Storz & Bickel Volcano ^®^, Tuttlingen, Germany), with participants inhaling equal amounts of each while lying in the MRI scanner. The treatments were vaporized at 225°C and the vapor was stored in a polythene bag equipped with a mouthpiece. Participants were instructed to place the mouthpiece to their lips, inhale deeply for 4 seconds, hold their breath for 10 seconds, and then exhale. Participants repeated this procedure until the balloon was empty. Participants were instructed to inhale the entire volume of the balloon within 5 minutes. Participants received two resting state scans at 15 min and 36 min after inhalation. Resting state scans were preceded by a psychomotor vigilance task and the collection of a blood sample and were directly followed by a rating of subjective high. Outcome parameters were averaged across successive measurements to yield a single value per individual in each treatment condition to serve as input for the connectome analysis. The current study was registered in the Netherlands trial register (NTR4897). A previous analysis of THC effects on seed-based functional connectivity within the mesocorticolimbic circuit employing the same data set has been published elsewhere ^11^.

Participants received a training day prior to the treatment conditions to become familiarized with the study procedures. Participants in the occasional users group were instructed to refrain from drug use, including cannabis, (≥7 days) and alcohol (≥24 hours) prior to their testing day; whereas participants in the chronic group were given the same instructions, however were allowed to use cannabis up until 24 hours prior to their testing day. Absence of drug and alcohol was assessed via a urine drug screen and a breath alcohol screen at the start of a test day. A pregnancy test was given if participants were female. If all tests were found to be negative (except for cannabis in the chronic group), participants were allowed to proceed.

The study was conducted according to the code of ethics on human experimentation established by the declaration of Helsinki (1964) and amended in Fortaleza (Brazil, October 2013) and was approved by the Academic Hospital and University’s Medical Ethics committee. All participants gave their written informed consent. A permit for obtaining, storing, and administering cannabis was obtained from the Dutch Drug Enforcement Administration.

### Subjective and behavioral measures

Sustained attention was assessed via the psychomotor vigilance task (PVT), a 5-minutes reaction-time task that measures the speed with which participants responds to a visual stimulus ^45^. The primary outcome measure of the task is the number of attentional lapses (reaction time >500 ms). Participants also rated their subjective high on visual analog scales (10 cm) on two consecutive time points after treatment administration, on a scale between 0 (not high at all) and 10 (extremely high). Both measures were conducted by the participants while in the scanner.

### Pharmacokinetic measures

Blood samples (8 mL) to determine cannabinoid concentrations (THC and metabolites OH-THC and THC-COOH) were taken at baseline and prior to resting state measures and analyzed according to a standardized procedure ^46^.

### Resting state functional connectivity

All participants underwent a resting state functional MRI. Images were acquired on a MAGNETOM 7T MR scanner. A total of 258 whole-brain EPI volumes were acquired at rest (TR = 1400 ms; TE = 21 ms; flip angle = 60°; oblique acquisition orientation; interleaved slice acquisition; 72 slices; slice thickness = 1.5 mm; voxel size = 1.5 × 1.5 × 1.5 mm). During scanning, participants were shown a black cross on a white background and were instructed to focus on the cross while attempting to clear their mind.

fMRI data were processed with an in-house developed pipeline based on Matlab and FSL, using state-of-the-art guidelines ^39,47,48^. These steps included: BOLD volume unwarping (FSL apply topup), slice timing correction (FSL slicetimer), realignment (FSL mcflirt), normalization to mode 1000, demeaning and linear detrending (Matlab detrend), regression (Matlab regress) of 18 signals: 3 translations, 3 rotations, and 3 tissue-based regressors (mean signal of wholebrain, white matter (WM) and cerebrospinal fluid (CSF)), as well as 9 corresponding derivatives (backwards difference; Matlab). We also kept track of the fMRI volumes that were highly influenced by head motion, by using three different metrics: 1) Frame Displacement (FD, in mm); 2) DVARS (D referring to temporal derivative of BOLD time courses, VARS referring to root mean square variance over voxels) (Power et al. 2014); 3) SD (standard deviation of the BOLD signal within brain voxels at every time-point). The FD and DVARS vectors (obtained with fsl_motion outliers) were used to detect outlier BOLD volumes with FD > 0.3 mm and standardized DVARS > 1.7. The SD vector obtained with Matlab was used to detect outlier BOLD volumes higher than 75 percentile + 1.5 of the interquartile range per FSL recommendation ^49^. A bandpass first-order Butterworth filter [0.009 Hz, 0.08 Hz] was applied to all BOLD timeseries at the voxel level (Matlab butter and filtfilt). As a final denoising step, the first three principal components of the BOLD signal in the WM and CSF tissue were regressed out of the gray matter (GM) signal (Matlab, pca and regress) at the voxel level.

A whole-brain data-driven functional parcellation based on 278 regions ^50^, was projected into each subject’s T1 space (FSL flirt 6dof and FSL flirt 12dof) and then into native EPI space of each subject. The voxelwise BOLD signals were averaged into the corresponding Shen brain regions, and then functional connectomes were computed as Pearson’s correlation between time series of all region pairs. Finally, the resulting functional connectomes (278 cortical and subcortical nodes) were ordered according to seven cortical resting state networks (RSNs) as proposed by Yeo et al. ^51^. These included the visual (VIS), somatomotor (SM), dorsal attention (DA), ventral attention (VA), limbic (L), frontoparietal (FP) and the default mode network (DMN). For completeness, we added two more networks: one composed of the subcortical and one for the cerebellar regions.

We explored functional connectome changes in a population of chronic and occasional cannabis users, using two different methodologies. The first, network-based statistics (NBS), is a connectome-wide analysis where hypothesis testing at each and every element of the connectivity matrix is performed. The second, connectivity independent component analysis (connICA), extracts independent subsystems of connectivity from the individual functional connectomes. These subsystems can then be associated with behavioral and demographic scores or scores related to treatment and cannabis use history.

### Network based statistics on functional connectomes of occasional and chronic users

The network-based statistic (NBS) is a popular network-specific approach to control the familywise error rate (FWER) when performing mass univariate testing on all connections in a functional or structural connectome ^43^, between two or more groups (e.g., in the case of this work, between occasional and chronic users). The NBS is used in settings where each connection is associated with a test statistic and corresponding p-value, and the goal is to identify groups of connections showing a significant effect while controlling the FWER. The approach is somewhat analogous to cluster-based approaches developed for performing inference on statistical parametric maps in human neuroimaging ^52,53^. Instead of identifying clusters of voxels in physical space, the NBS identifies connected subnetworks in topological space. The size of a subnetwork is most typically measured by the number of edges that it comprises. A summary description of the NBS workflow is given in Figure 1A.

**Fig. 1.**
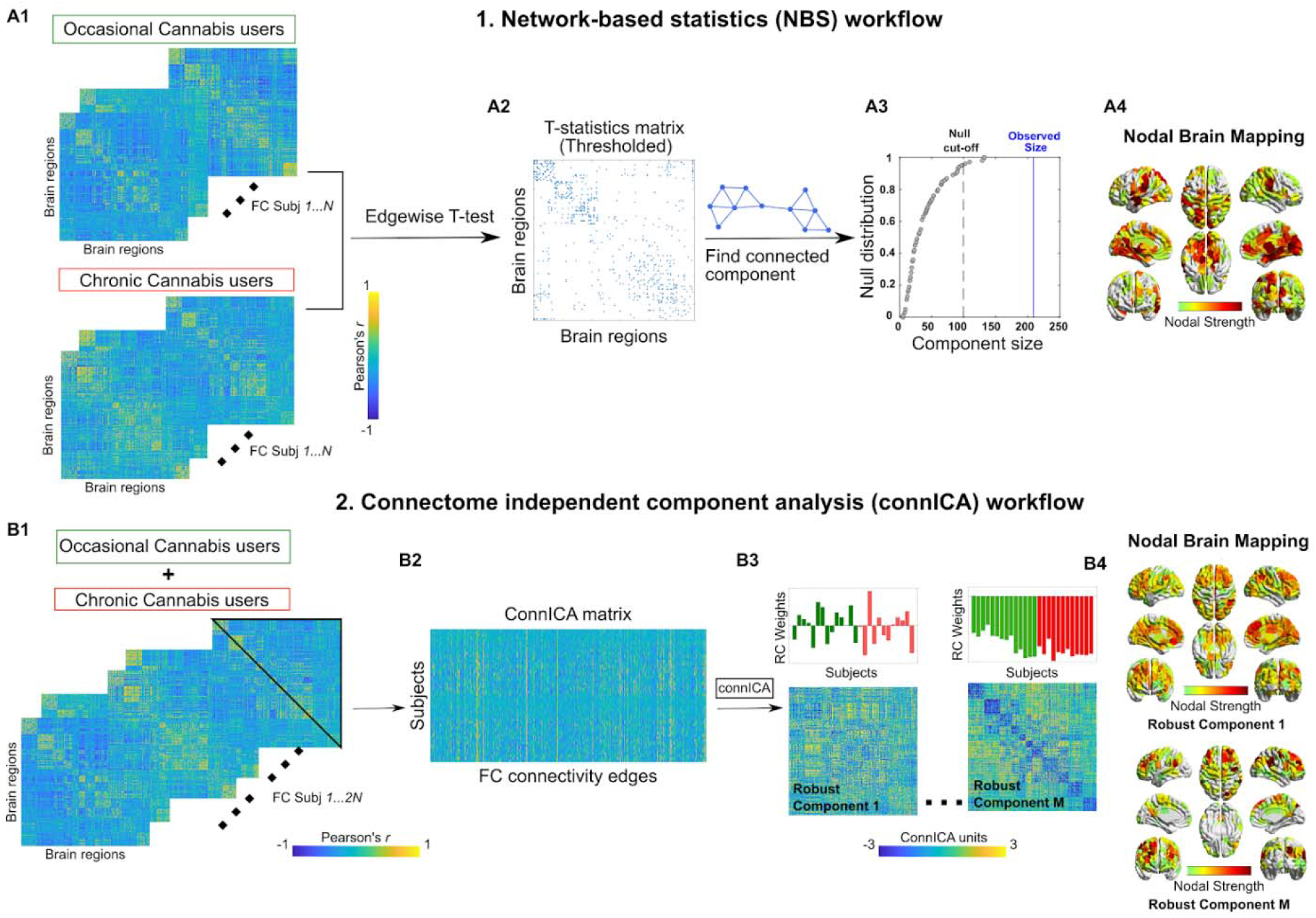
Workflows of NBS (upper panel) and connICA (lower panel). **A1)** NBS starts from two groups of functional connectomes, in this case from occasional and chronic cannabis users. **A2)** A t-test is performed edgewise resulting in a matrix of T-statistic values. This matrix is then thresholded to yield a binarized T-statistic matrix. **A3)** The giant component (largest connected component) of the T-statistic matrix is computed and its size compared with the ones obtained by doing T-statistic by randomly shuffling occasional and chronic users’ functional connectivity. The red line indicates the cut-off for declaring a component size as statistically significant (α < 0:05). The blue line shows the observed size of the component for T-matrix in A2. **A4)** Brain render of the nodal strength of the regions most involved in the T-statistics. **Bottom)** connICA ^39^ workflow. **B1)** The upper triangular elements of each individual functional connectivity matrix (from both occasional and chronic users) are added to **B2)** a matrix whose rows indicate subjects, and columns subjects’ vectorized whole-brain functional connectomes. **B3)** The ICA algorithm extracts M independent components (i.e., patterns) associated with the whole population and their relative weight in each subject. Functional connectivity, expressed as Pearson’s correlation coefficient values for individual FC matrices. The bar plots above the FC components indicate the individual prominence or amount (i.e. weights) of each extracted FC independent component in the original single-subject FC on the left side of the figure. They are color-coded according to the group membership (i.e. chronic or occasional weights). **B4)** Brain render of the nodal strength of the regions most involved in a specific connICA robust component.

Briefly, NBS independently computed a univariate test statistic (e.g. t-statistic) between the groups for each and every connection ^54^. The result is a matrix of test statistic values with the same dimensions as the connectome. We then threshold the test statistic matrix to keep only the significant edges. This thresholded matrix is then connected to components, that is subnetworks of edges showing a common statistical effect of interest (Fig. 1A2). The size of each component is stored. The size of a connected component can be measured as the number of edges it comprises. After the sizes of the observed components are computed, permutation testing is used to estimate a corrected p-value for the size of each observed component, by randomly shuffling the labels assigned to each network so that the “random” groups comprise a mixture of actual occasional and chronic cannabis users. The analysis is then repeated and the size of the largest component is stored. We repeat this procedure many times to generate an empirical null distribution of maximal component size. The corrected p-value for a component of size m is then given by the proportion of permutations for which the largest component is equal to or greater in size than m (Fig. 1A3).

Here we used NBS to investigate whole-brain connectivity changes in functional connectomes of chronic and occasional users groups, during the placebo and the THC condition independently. In order to detect mean differences between functional connectome edges, we used double-sided t-test (thresholded at p<0.01, i.e. T score = 3.1). The cutoff alpha for the giant component size was set to 0.05.

### Connectivity independent component analysis (connICA) on functional connectomes of occasional and chronic cannabis users

ConnICA is a novel data-driven methodology that applies independent component analysis ^39^ to extract independent connectivity patterns from individual functional connectomes. Output of connICA includes: i) an “FC-component” representing an independent pattern of functional connectivity present across participants, and ii) each participant’s weight, quantifying the (signed) component strength or prominence in each individual FC matrix. A summary description of the connICA workflow is given in Figure 1B.

After the initial NBS analysis where we aimed at looking into widespread changes in chronic and occasional users’ functional connectomes, we next applied connICA to zoom into the independent “components of interest” that significantly differentiated chronic and occasional users, as well as those that were associated with the acute intoxication effect of THC. Note that in this case we used all the functional connectomes from the two groups and the two conditions, to maintain the “blind data-driven” spirit of the connICA framework.

Given the non-deterministic nature of the ICA decomposition into components ^55,56^, multiple ICA runs are required to select the most robust outcomes ^39,55^. As in previous work, we accounted for this by evaluating the robustness of the components (“FC-traits” in ^39^) over 100 FastICA runs. The FC-component was considered robust when it appeared in at least 75% of the runs, as defined by a correlation of 0.75 or higher across runs ^39^. Before running the connICA algorithm, we applied Principal Component Analysis ^57^ to perform noise filtering and dimensionality reduction, as recommended by work in machine learning ^58^ and neuroimaging communities ^59,60^. After this PCA-based preprocessing, we estimated the number of independent components ^39,61^. The two parameters of percent retained variance from PCA and number of independent components were broadly explored to find the optimal combination. For each block, we examined percent variance retained after PCA in the range [75%, 100%], in steps of 5%. Similarly, we evaluated the number of ICA components in the range [5, 25], in steps of 1. Considering our a-priori hypothesis, we aimed for the range of parameters where we had the greatest number of robust components, while preserving most of the information from the data (minimal PCA reduction). As depicted in Fig. S1, the optimal choice of these two parameters was 95% retained variance in PCA and 20 independent components.

After connICA extraction of the most robust connectivity patterns from the dataset, we performed a multi-way analysis of variance model (MATLAB anovan), in order to account for the interactions between the different factors, including the repeated measures (i.e. participants’ connectomes appearing repeatedly, in different conditions) effects. The predictors included in the ANOVA model were: Drug condition (THC or placebo); cannabis user group (occasional or chronic cannabis users), gender, age, rating of subjective high and number of lapses of attention as assessed in the psychomotor vigilance task.

Finally, we evaluated and reported the connICA components where the Anova model showed significant associations for the predictors of interest, after Bonferroni correction for multiple comparisons across the number of components tested (threshold was set to p<0.01, Bonferroni corrected).

### Statistics on subjective and behavioral measures

A mixed-model analysis was performed consisting of the within-subject factors treatment (THC and placebo) and the between-subject factor of group (occasional or chronic) on mean subjective high ratings and number of attentional lapses averaged across two successive time points. The alpha criterion level of significance was set at p = 0.05.

## Results

### Subjective and behavioral data

Mixed-model analyses of variance (ANOVA) yielded a significant main effect of Treatment on ratings of subjective high [F(1,23) = 40.26, p = <.001, *ηp*^2^ = .641] and the number of lapses of attention [F(1,23) = 4.71, p = .041 *ηp*^2^ = .169], indicating that subjective high and lapses of attention were higher in the THC condition as compared to the placebo condition. Separate contrasts revealed that THC increased lapses of attention primarily in occasional users [F(1,12) = 5.39, p = .039, *ηp*^2^ = .310], but not in chronic users. The number of lapses of attention were also significantly higher in occasional users as compared to chronic cannabis users [F(1,23) = 9.21, p = .006, *ηp*^2^ = .238]. Mean (SE) subjective ratings of high and lapses of attention are given in Table S2.

### THC concentrations in serum

Mean (SE) concentrations of THC, 11-OH-THC, and THC-COOH in serum are given in Table S3. As expected from previous experience ^62^, THC [F(2,32) = 6.29, p = .023] and THC-COOH [F(2,34) = 6.29, p = .031] levels were higher in chronic users as compared to occasional users even though they received the same dose.

### Network based statistics

We performed NBS T-statistic between the two cannabis user groups, when they were in the placebo condition and while they were under the influence of THC (Figure 2). Interestingly, NBS showed a significant broad difference in the functional connectome between the 2 users groups while under the influence of THC (T-matrix thresholded at T Score of 3.1, which equals p<0.01, Fig. 2A-D). The functional connectome revealed (for T-test chronic>occasional) increments in functional connectivity within most Yeo-Networks (Figures 2B-D). T-tests comparing occasional>chronic mainly showed significant changes in network connectivity between DMN and dorsal and ventral attentional networks (Fig 2B-D). Group differences were less apparent during the placebo condition. Only the T-test chronic>occasional revealed a significant difference in the sensory-motor network. NBS did not reveal any significant differences when comparing THC vs placebo conditions in each of the cannabis user groups (see Figure S2).

**Fig. 2.**
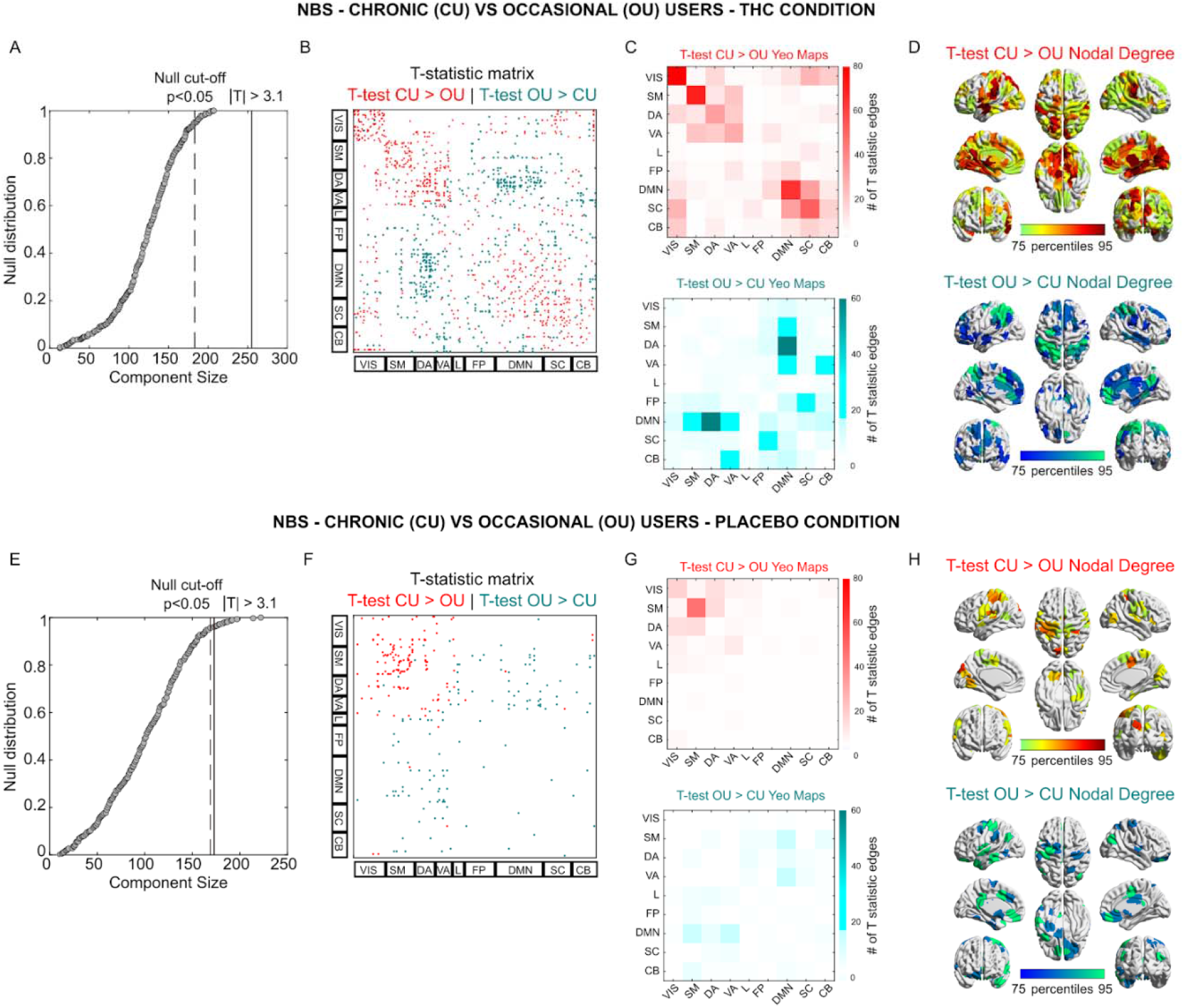
Network-based analysis of functional connectome changes between occasional and chronic cannabis users in each treatment condition. **B-F)** T-statistic matrix threshold at |T|>3.1 (which corresponds to double-sided p< 0.01) for edgewise group differences between chronic users (CU) and occasional users (OU), during THC (B) or Placebo (F) condition. The brain regions are ordered according to the resting-state network organization proposed by Yeo et al (REF). **A-E)** Giant component size extracted from the T-statistic matrix on the observed effect (Red for CU>OU; Teal for OU>CU) and the null distribution obtained by randomly shuffling chronic and occasional functional connectomes. **C-G)** Mapping of the significant edges into the 7 functional networks by Yeo (REF), with the addition of subcortical and cerebellar networks, for the THC (C) and Placebo condition (G). **D-H)** Brain renders reporting the sum of the significant edges where CU>0U and OU>CU, during THC (D) and Placebo (H) condition.

### ConnICA

ConnICA allowed us to identify spatial functional connectome patterns that were related to chronic use of cannabis and to the acute state of THC intoxication. ConnICA extracted two components of interest. The first was associated with differences between the cannabis user groups across treatment conditions (Figure 3). The second was associated with differences in subjective high during cannabis and placebo, across the two cannabis user groups (Figure 4). The first component identified greater functional connectivity in chronic cannabis users as compared to occasional cannabis users in all networks (anova p < 0.01, Bonferroni corrected for multiple comparisons across connICA components). This finding appears broadly in line with the between cannabis user group differences that was obtained with NBS. The second component identified a functional connectivity pattern that was associated with the subjective state of cannabis intoxication as rated on visual analog scales of subjective high (anova p < 0.01, Bonferroni corrected for multiple comparisons across connICA components). The pattern consisted of an increase in functional connectivity between the default mode network and the ventral attentional network, and decreased functional connectivity between the subcortical network and the dorsal attentional network and between the cerebellum and the limbic network. Age and gender were not associated with a connICA pattern. A functional connectivity pattern associated with attentional performance did not survive the correction for multiple comparisons.

**Fig. 3.**
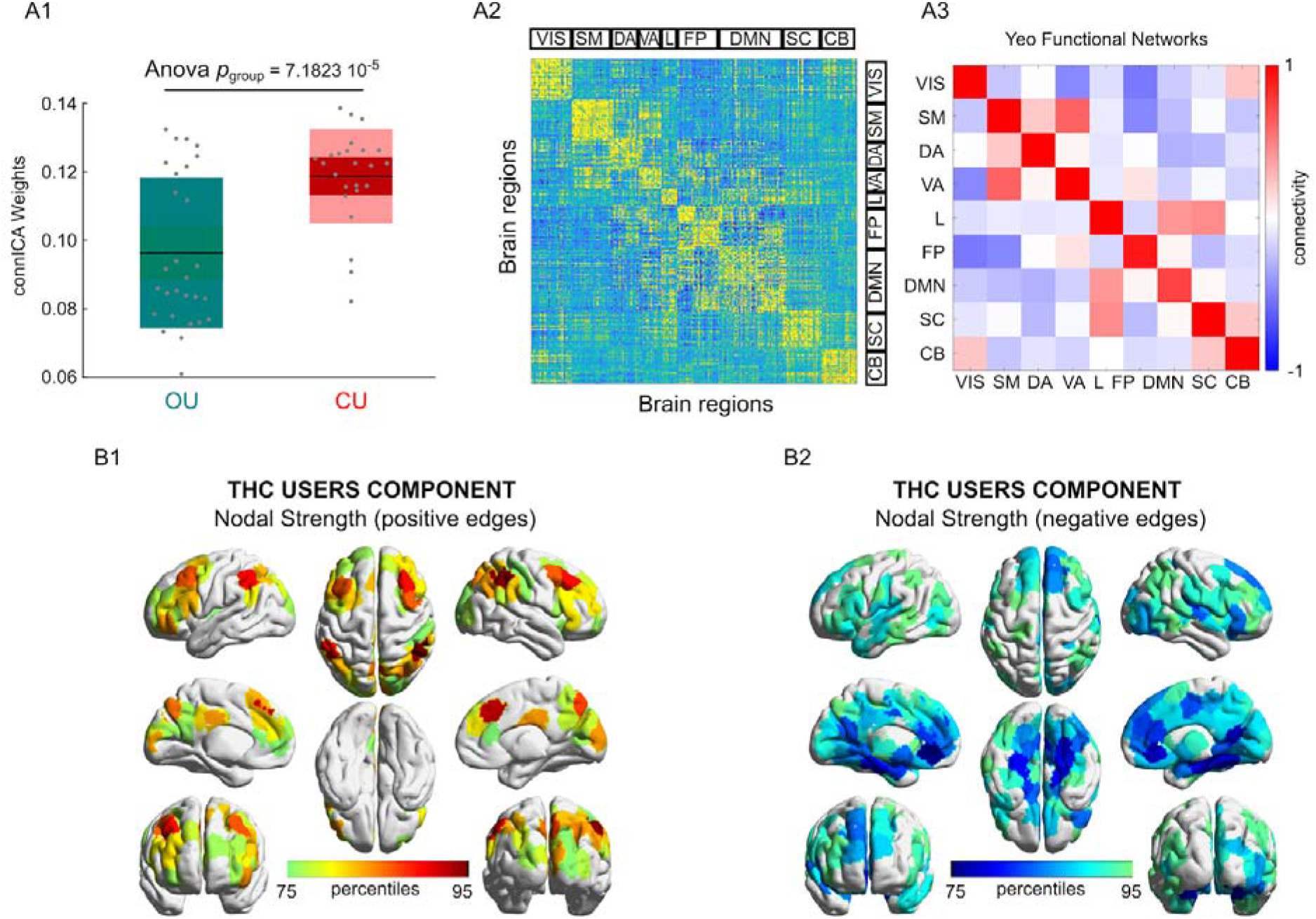
The cannabis user group-differentiating connICA component. **A1)** Group differences in the individual subject weights associated with the connICA component (anova test, p < 0.01 Bonferroni corrected across the 20 robust components, see Methods); **A2)** The functional connectivity component extracted by connICA. Pairwise associations between brain regions are ordered by resting-state functional networks as proposed by Yeo et al. ^51^. **A3)** For clarity, the same component, depicted after averaging across functional networks, shows prominent connectivity within the main functional network areas. **B1-B2)** Nodal strength (sum over columns of A1, including only: B1) positive edges or B2) negative edges) of the top 25% regions involved in the identified group-differentiating component.

**Fig. 4.**
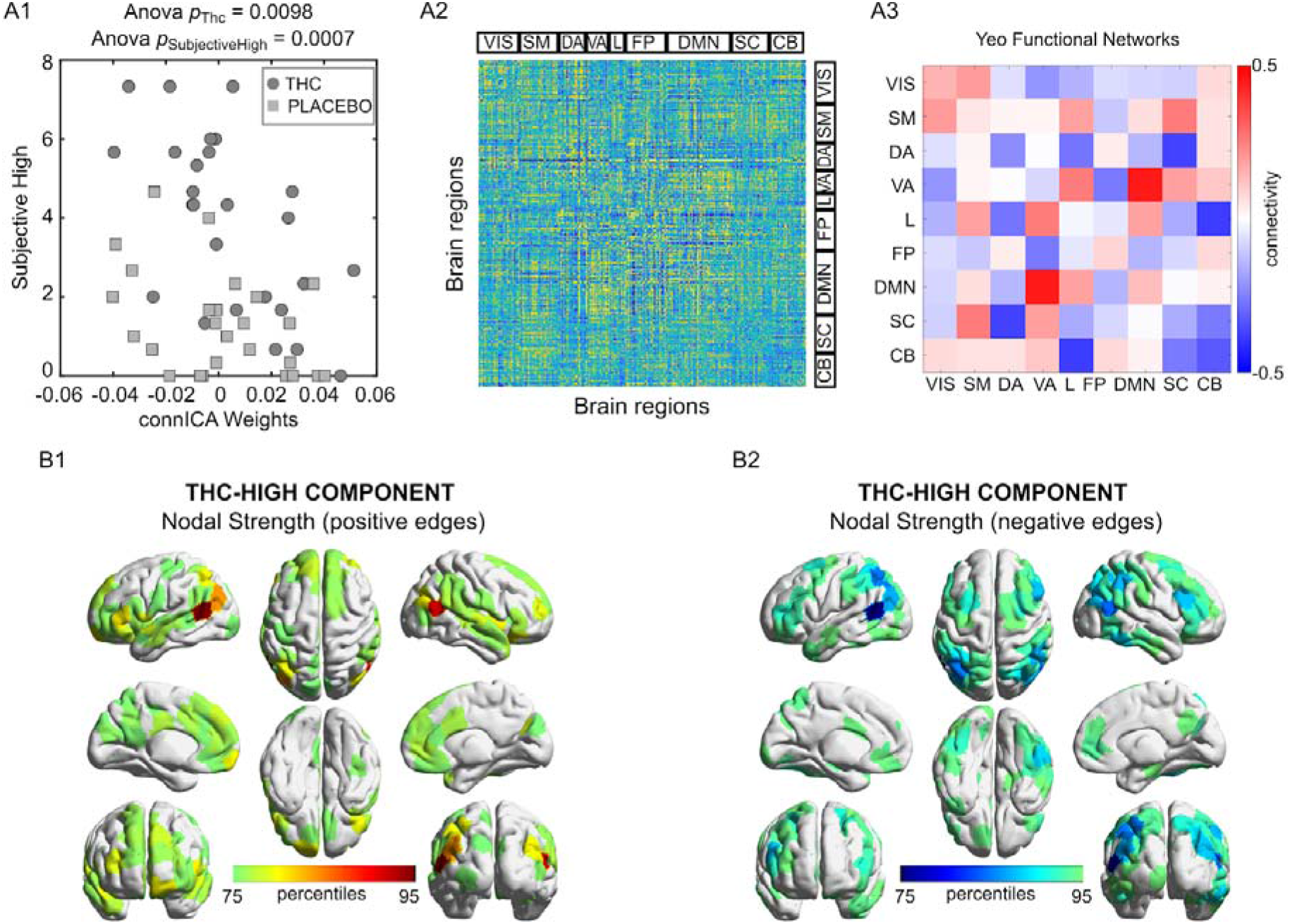
THC induced feeling of subjective high connICA component. **A1)** Drug differences in the individual subject weights associated with the connICA component (anova test, p < 0.01 Bonferroni corrected across the 20 robust components, see Methods) in association to subjective high in each drug condition (circles, THC; squares, placebo). The connICA component and subjective high ratings are strongly correlated (see Methods for details). **A2)** The functional connectivity component extracted by connICA. Pairwise associations between brain regions are ordered by resting-state functional networks as proposed by Yeo et al. ^51^. **A3)** For clarity, the same component, depicted after averaging across functional networks, shows prominent connectivity within the DMN and attentional areas. **B1-B2)** Nodal strength (sum over columns of A1, including only: B1) positive edges or B2) negative edges) of the top 25% regions involved in the identified THC-differentiating component.

## Discussion

The present study aimed to determine the effects of acute and chronic cannabis use on the whole brain connectome using two previously established methodologies, i.e. NBS and connICA, which provided consistent and complementary results. Both NBS and connICA revealed strong increments in functional connectivity within the major brain networks in chronic cannabis users as compared to occasional users, suggesting a state of hyperconnectivity. Likewise, both methodologies showed that increments *within* network functional connectivity in chronic cannabis users were paralleled by decrements in connectivity *between* networks. The impact of cannabis use history on the human connectome appeared more prominent when using the connICA methodology that allows extraction of spatial patterns of connectivity associated with clinical characteristics. Apart from a state of hyperconnectivity in chronic cannabis users, the connICA methodology also extracted a distinct spatial connectivity pattern associated with the feeling of subjective high during cannabis intoxication across both users groups.

Acute cannabis intoxication produced a select, spatial pattern in functional connectivity that was strongly associated with the feeling of subjective high in both user groups. Most prominent were decrements in functional connectivity between the subcortical and the dorsal attention network, and between the limbic and cerebellar networks that suggest a reduction of top-down attention control ^63^ and motor coordination ^64^ during cannabis intoxication. These findings of reduced connectivity are largely in line with a series of studies that predominantly reported hypoconnectivity within brain circuits as a primary response during cannabis intoxication ^10,11,27^. A parallel increment in functional connectivity however was apparent between the default mode network and the ventral attention network. The ventral attention network is assumed to be involved in stimulus-driven shifts of attention ^63^ and its increased connectivity with the self-centered default mode network suggest an increased internalization of attention directed at monitoring of mind wandering or spontaneous cognition ^65^ during cannabis intoxication. As such, the spatial functional connectivity pattern identified by connICA reflects some of the core elements of cannabis intoxication including the feeling of subjective high and alterations in attentional and motor function ^66^.

NBS revealed hyperconnectivity within a number of brain networks of chronic cannabis users. ConnICA subsequently identified hyperconnectivity within all brain networks as a significant pattern that distinguished chronic cannabis users from occasional cannabis users. In addition, moderate increments in functional connectivity in chronic cannabis users were also observed between a select number of networks: i.e. between the limbic network and the DMN and subcortical brain areas, and between attentional networks and the somatomotor network. This broad pattern of hyperconnectivity was also paralleled by moderate reductions of functional connectivity between remaining network edges. The prime finding of hyperconnectivity across the whole brain connectome of chronic cannabis users adds to findings of previous imaging studies showing increased functional connectivity in subcortical and frontal regions ^33,37^ or between those regions ^34–36^. The present study, however, suggests that functional hyperconnectivity in chronic cannabis users is not restricted to local brain circuits but can be observed across the entire whole brain connectome.

The widespread pattern of hyperconnectivity in the human connectome of chronic cannabis users can be interpreted in various ways. It may indicate a general downregulation of CB1 receptors that are expressed across the entire central nervous system ^67^. A number of studies have shown a global reduction in CB1 receptor availability of 10-20% across the brain of chronic cannabis users ^21–23^ that might be expected to produce an operational disruption in neural signaling within the neurotransmitter circuits in which they are expressed ^3,4^. Previous studies have suggested that hyperconnectivity observed in resting state fMRI is a common response to neurological disruption that may be differentially observable across the entire brain ^68,69^. In the context of stimulant drug abuse, development of hyperconnectivity within brain circuits has also been related to countervailing resilience systems implicated in behavioral regulation and compensation ^70^. The finding that acute (i.e. hypoconnectivity) and chronic effects (hyperconnectivity) of cannabis on brain circuits are largely opposite may support the notion that chronic effects produced by cannabis might reflect a compensatory response of the brain to repeated cannabis use and thus stimulations of CB1 receptors. That also seems to be in line with the finding that chronic users of cannabis can develop (partial) tolerance to acutely impairing effects of cannabis, presumably as a consequence of adaptive CB1 receptor downregulation ^12^.

Interestingly, none of the spatial patterns that were identified by connICA were associated with attentional performance differences between cannabis user groups or between treatments even though these were evident from performance data. Effects sizes of differences in attentional performance however were relatively small and, in case of a treatment effect, limited to occasional users. This may have hindered a clear-cut attribution of attentional changes to a functional connectivity pattern across the cannabis user groups, particularly after corrections for multiple comparisons. Alternatively, changes in attentional performance may also be driven by local changes in a confined brain circuit that may go undetected in a spatial analysis of whole-brain network patterns. Previous analyses have indeed linked THC induced impairment of attention to a reduction in functional connectivity within (sub)cortical areas of the reward circuit of occasional cannabis users ^10^, but not in chronic cannabis users ^11^. In both cases, seed-based connectivity analyses were used to test the particular hypothesis that THC-induced increment in dopamine release to the nucleus accumbens would alter its connectivity with neural structures in the reward circuit. Such lower-level, circuit-specific changes may be harder to identify with higher-level aggregation methods as employed in the present study.

Overall, the present study suggests an important utility for whole-brain network approaches in the identification and separation of acute and chronic effects of cannabis on the functional brain connectome. Incorporation of cross-network dynamics might identify neurobiological features or phenotype characteristics of impaired and adaptive behaviors that might arise during acute and chronic use of cannabis. Such models might provide unique insights into the emergence and maturation of distinct functional networks in users that progress from acute to chronic cannabis use, and into temporal alterations in network dynamics that underlie the development of pathological states ^71^, e.g. cannabis use disorder or cannabis-induced psychosis, in a subset of users. In general, approaches that focus on large-scale brain organization or distributed brain circuits are well equipped to capture the complexity of brain function ^72^ and subsequently may also be best suited to assess alteration in functional brain dynamics following cannabis use.

In conclusion, we have shown with two separate methodologies that whole-brain functional connectomes can distinguish occasional from chronic cannabis users and identify acute cannabis intoxication, linking brain network dynamics with cannabis-induced long and short-term changes. This work is relevant for probing the neurobiological basis of behavioral function and dysfunction related to cannabis use, and its associated brain network dynamics, as well as to substance use in general.

## Supplementary Information

**Fig. S1.**
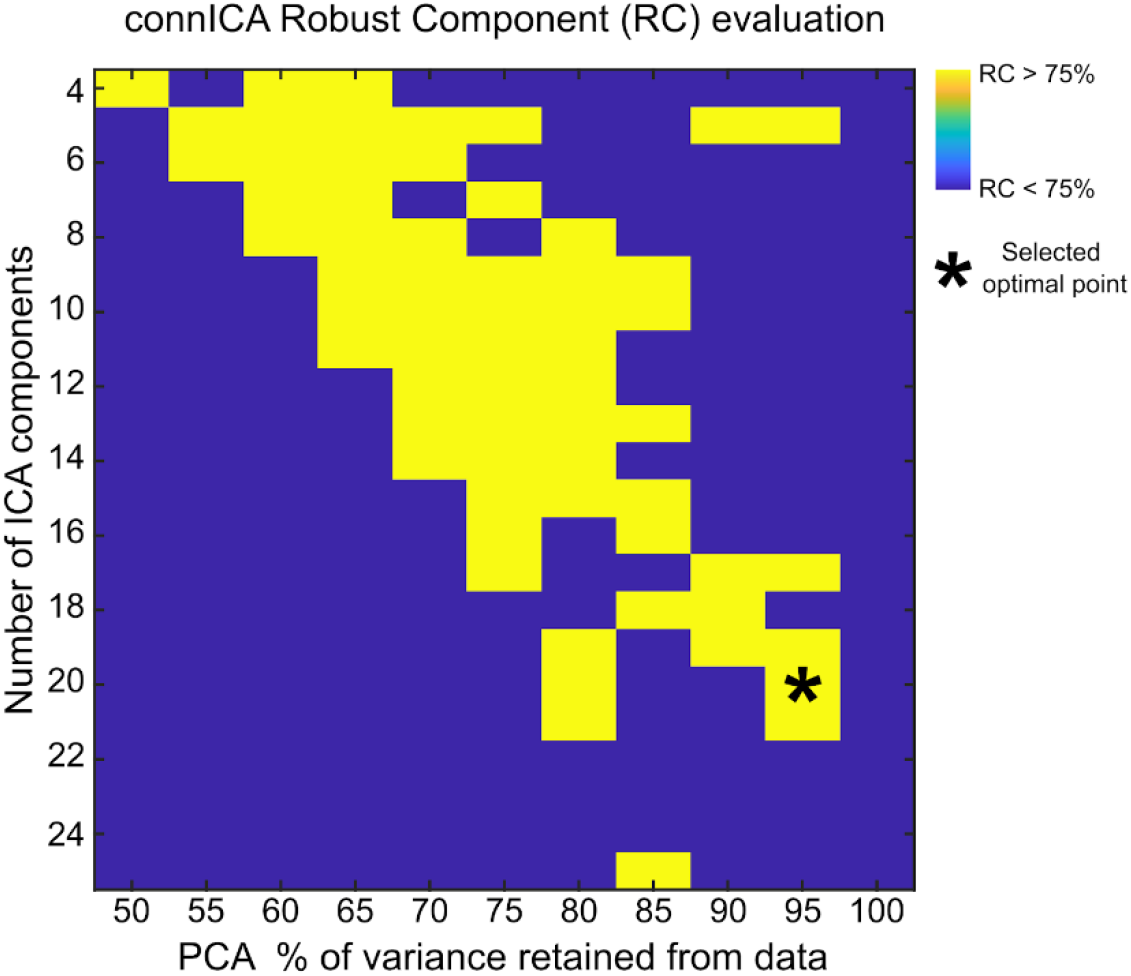
Exploration of optimal connICA parameters. The two free parameters of the connICA methodology ^39^, i.e. the number of ICA components and the percentage of variance retained, were explored to maximize: 1) the number of robust components and, 2) the percentage of variance retained from the data. The optimal point at 95% retained variance in PCA and 20 independent components is indicated by an asterisk.

**Fig. S2.**
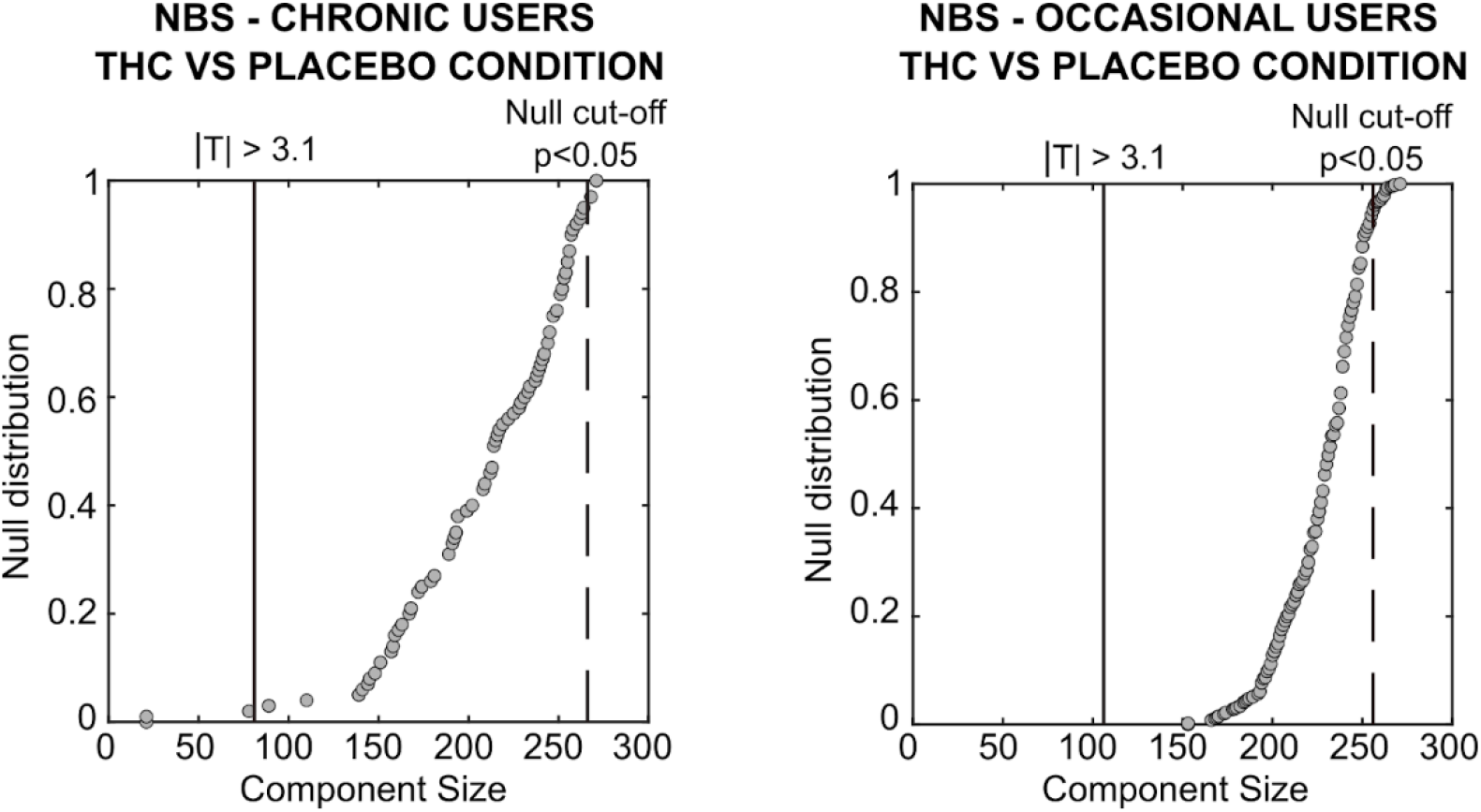
NBS analysis of THC vs placebo effects in each cannabis user group. Giant component size extracted from the T-statistic matrix obtained when comparing THC vs placebo conditions within the cannabis user groups. Note how the effect does not survive the cut-off threshold obtained by random shuffling the drug labels, in both groups

**Table S1.**
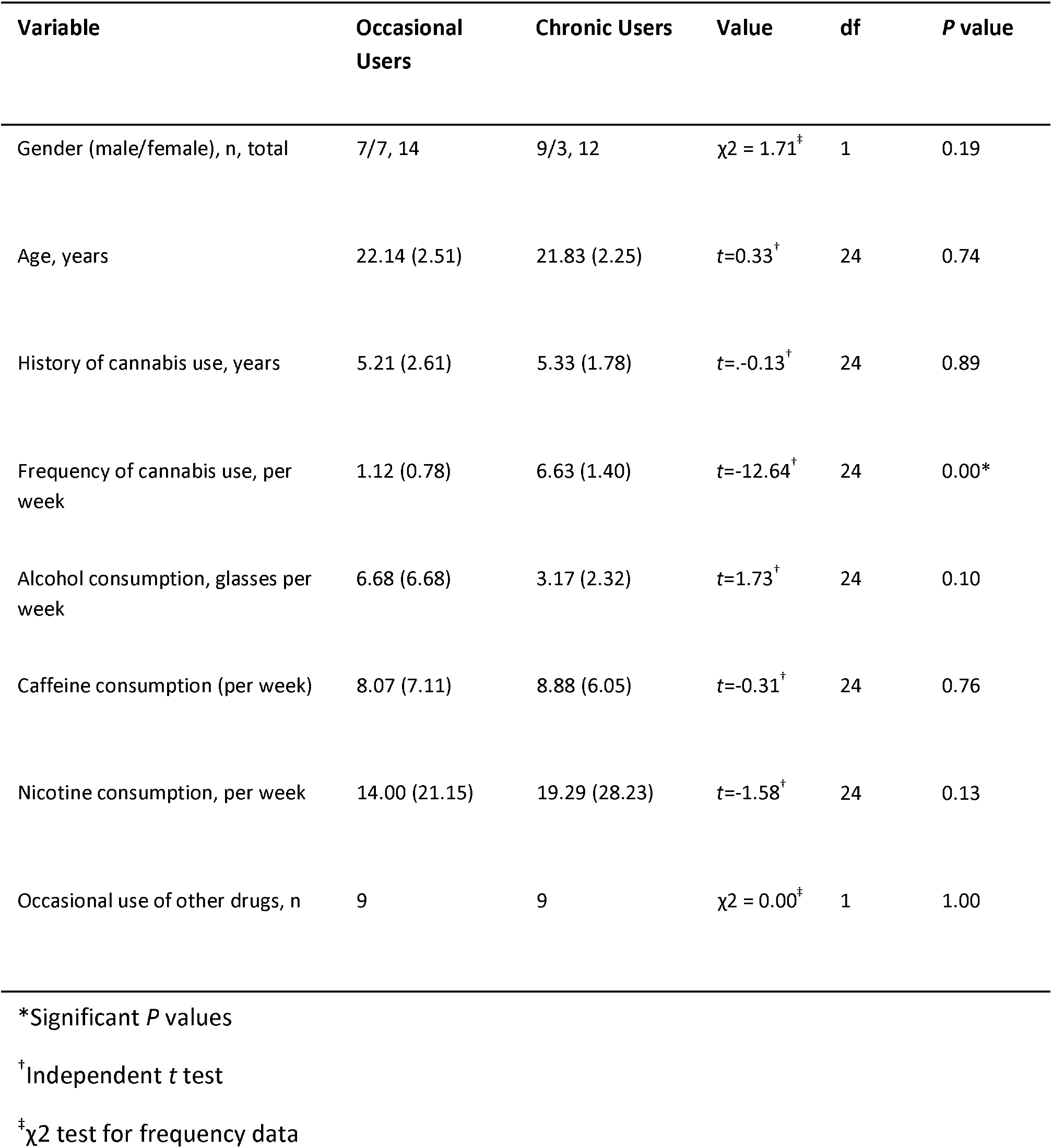
Mean subject characteristics (SD) and history of drug use for occasional and chronic cannabis users who completed the study (N=26).

**Table S2.** Mean (SE) subjective high, lapses of attention averaged over two time points in each treatment condition. OU=occasional users; CU=chronic users.

|  | Subjective High (cm) |  |  | Lapses of attention (#) |  |  |
| --- | --- | --- | --- | --- | --- | --- |
|  | Post-resting<br>state 1<br>(20 min) | Post-resting<br>state 2<br>(42 min) | average | Pre-resting<br>state 1<br>(1 min) | Pre-resting<br>state 2<br>(22 min) | Average |
| **OU** |  |  |  |  |  |  |
| Placebo | 1.21 (.38) | 0.92 (.37) | 1.07 (.31) | 3.50 (.81) | 5.35 (1.51) | 4.42 (1.08) |
| THC | 5.36 (.70) | 3.92 (.61) | 4.73 (.61) | 5.0 (1.26) | 8.07 (1.78) | 6.15 (1.38) |
| **CU** |  |  |  |  |  |  |
| Placebo | 2.50 (.62) | 2.08 (.62) | 2.29 (.57) | 1.25 (.30) | 1.08 (.31) | 1.16 (.23) |
| THC | 4.83 (.75) | 4.08 (.73) | 4.45 (.71) | 1.83 (.34) | 1.91 (.62) | 1.87 (.35) |

**Table S3.**
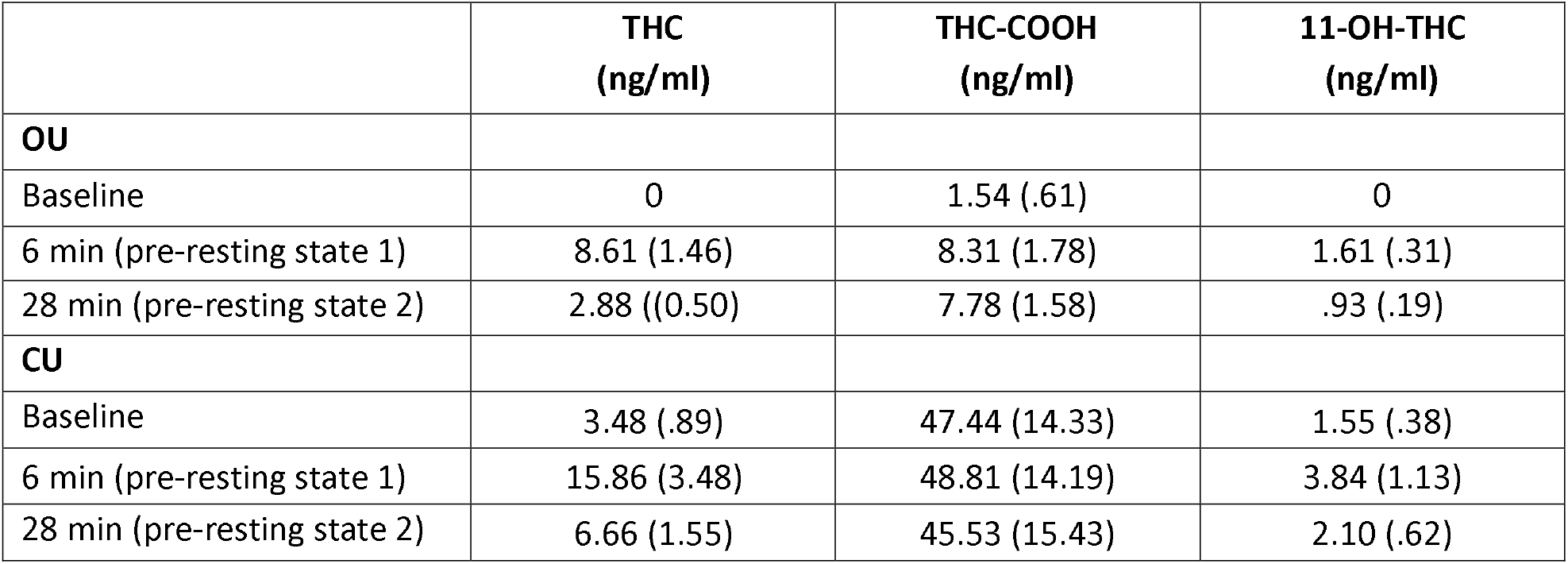
Mean (SE) THC, THC-COOH and 11-OH-COOH concentrations at baseline and two time points prior to resting state fMRI in THC condition. OU=occasional users; CU=chronic users.

|  | THC<br>(ng/ml) | THC-COOH<br>(ng/ml) | 11-OH-THC<br>(ng/ml) |
| --- | --- | --- | --- |
| **OU** |  |  |  |
| Baseline | 0 | 1.54 (.61) | 0 |
| 6 min (pre-resting state 1) | 8.61 (1.46) | 8.31 (1.78) | 1.61 (.31) |
| 28 min (pre-resting state 2) | 2.88 ((0.50) | 7.78 (1.58) | .93 (.19) |
| **CU** |  |  |  |
| Baseline | 3.48 (.89) | 47.44 (14.33) | 1.55 (.38) |
| 6 min (pre-resting state 1) | 15.86 (3.48) | 48.81 (14.19) | 3.84 (1.13) |
| 28 min (pre-resting state 2) | 6.66 (1.55) | 45.53 (15.43) | 2.10 (.62) |

